# Increased p53 signaling impairs neural differentiation causing HUWE1-promoted intellectual disabilities

**DOI:** 10.1101/2020.05.10.087296

**Authors:** Rossana Aprigliano, Stefano Bradamante, Boris Mihaljevic, Wei Wang, Sarah L. Fordyce Martin, Diana L. Bordin, Matthias Bosshard, Nicola P. Montaldo, Yunhui Peng, Emil Alexov, Cindy Skinner, Nina-Beate Liabakk, Magnar Bjørås, Charles E. Schwartz, Barbara van Loon

## Abstract

Essential E3 ubiquitin ligase HUWE1 (HECT, UBA and WWE domain containing 1) regulates key factors, as p53. Mutations in *HUWE1* have been associated with neurodevelopmental X-linked intellectual disabilities (XLIDs), however the pathomechanism at the onset of heterogenous XLIDs remains unknown. In this work, we identify p53 signaling as the process hyperactivated in lymphoblastoid cells from patients with HUWE1-promoted XLIDs. The hiPSCs-based modeling of the severe HUWE1-promoted XLID, the Juberg Marsidi syndrome (JMS), reviled majorly impaired neural differentiation, accompanied by increased p53 signaling. The impaired differentiation results in loss of cortical patterning and overall undergrowth of XLID JMS patient-specific cerebral organoids, thus closely recapitulating key symptoms, as microcephaly. Importantly, the neurodevelopmental potential of JMS hiPSCs is successfully rescued by restoring p53 signaling, upon reduction of p53 levels. In summary, our findings indicate that increased p53 signaling leads to impaired neural differentiation and is the common cause of neurodevelopmental HUWE1-promoted XLIDs.

## INTRODUCTION

Neurodevelopment is a complex, finely regulated process. In this process, ubiquitination plays a fundamental role by controlling key factors that act in cell proliferation, differentiation, death and migration (1). The specificity of this posttranslational multistep mechanism is determined by E3 ubiquitin ligases that transfer ubiquitin to the substrate protein. Alterations in E3 ubiquitin ligases were associated with different neurological diseases. Mutations in E3 ligase *HUWE1* (HECT, UBA and WWE domain containing 1; also known as Mule, ARF-BP1, E3Histone, UREB1, HectH9 and LASU1) cause neurodevelopmental X-linked intellectual disability (XLID) syndromes (2-6). Clinical findings associated with HUWE1-promoted XLID syndromes are heterogenous and vary in severity, ranging from dysmorphic facial features and mild ID, to extreme ID and early death. Juberg-Marsidi syndrome (JMS) is one of the most severe forms of XLID, characterized by acute learning disability, generalized undergrowth, microcephaly, seizures and reduced life span (4). We recently showed that XLID JMS is caused by p.G4310R HUWE1 mutation and is accompanied by p53 accumulation (4). If and to which extent p53 accumulation contributes to JMS is unclear. Taken together, while different *HUWE1* mutations clearly cause XLID syndromes, the underlying pathomechanism through which *HUWE1* mutations contribute to onset of XLID remains unknown.

As E3 ubiquitin ligase Huwe1 was demonstrated to have vital functions in neurodevelopment, and its loss leads to lethality in mice (7, 8). Huwe1 impacts laminar pattering, by ensuring timely cell cycle exit and differentiation of neural progenitor cells (NPCs) in the cerebral cortex, and of neuronal and glia progenitors in the cerebellum (7, 9, 10). We suggested that similarly to mouse progenitors, HUWE1 has an important role in human NPCs (11). HUWE1 mediates its vital roles by regulating activity and stability of key cellular factors, as p53. In unstressed cells HUWE1 was suggested to be one of the key E3 ligases responsible for p53 ubiquitination and degradation (12). Transcription factor p53 controls expression of genes involved in cell cycle, apoptosis and differentiation. While p53 has been extensively studied as tumor suppressor whose loss promotes cancer, several lines of evidence suggested that altered p53 activity contributes to the developmental defects in different human genetic syndromes (13). By regulating the balance between apoptosis, proliferation and differentiation, p53 is suggested to play an important role in establishment of brain cytoarchitecture. Lack of p53 was shown to promote expansion of NPCs and alter their differentiation (14, 15), while p53 increase in neurons triggered developmental programmed cell-death (16-18). Though both HUWE1 and p53 play an important role in regulation of proliferation and differentiation, the extent to which these two factors co-operate in ensuring unperturbed neurodevelopment remains largely unknown.

In this work, we showed that increased p53 signaling is the common feature of the HUWE1-promoted XLID syndromes. Comparison of transcriptomes from XLIDs and healthy age-matched control cells identified a set of common deferentially expressed genes, which most significantly belong to the p53 signaling pathway. To explore the functional importance of deregulated p53 signaling, we focused on one of the most severe XLID forms the JMS caused by mutated HUWE1 p.G4310R. The increased p53 signaling in JMS patient-derived lymphoblastoid cells (LCLs) resulted in impaired cell cycle progression, reduced proliferation and increased apoptosis. Interestingly, the hiPSC-based JMS modeling revealed dramatically impaired neural differentiation capacity, accompanied by hyperactivated p53 signaling in JMS neural cells. The detected defects were reflected in large failure of JMS cerebral organoids to develop. Those JMS organoids that did develop had reduced size and severely impaired cortical patterning, closely recapitulating the microcephaly and undergrowth, key JMS symptoms. The observed neurodevelopmental defects are directly caused by elevated p53 activity, since p53 knockdown efficiently rescued neurodevelopmental potential of JMS hiPSCs. Taken together, our findings identify increased p53 signaling as the common pathomechanism at the onset of HUWE1-promoted XLIDs and suggest a crucial role of HUWE1-p53 pathway during human neurodevelopment.

## RESULTS

### Increased p53 signaling is a common process deregulated in XLID patients with mutated *HUWE1*

Mutations in *HUWE1* cause heterogenous neurodevelopmental XLID syndromes. To determine the common process that underlies *HUWE1*-promoted XLID, and since HUWE1 regulates multiple transcription factors important for neurodevelopment (19), we performed the RNA sequencing analysis. The transcriptomes from five XLID patient-derived LCLs, harboring different *HUWE1* mutations were compared with the transcriptomes from four healthy age- and sex-matched LCLs. This led to identification of 277 common differentially expressed genes (DEGs) in all XLID LCLs, at ≥1.5-fold change and an FDR ≤ 0.3. To determine the process commonly deregulated in all XLID LCLs, KEGG pathway analysis was performed next. The analysis of 277 DEGs revealed p53 signaling as the most significantly enriched pathway (Figure 1A). Interestingly, nearly all DEGs regulated by p53 were upregulated in XLID patient LCLs (Figure 1B), with the exception of *RRM2*, a ribonucleotide reductase gene known to be suppressed by p53 (20). These results suggest increased p53 signaling in all HUWE1-promoted XLID patients. This was further validated by RT-qPCR analysis of PUMA, GADD45α and p21 mRNA levels, the well-known targets of p53 (21). In accordance with RNA sequencing results, mRNA levels of these targets were significantly increased in XLID, when compared to the healthy individual cells (Figure 1C – 1E). Taken together, these data indicate that increased p53 activity, and subsequent deregulation of p53 targets, are the common features of differentiated XLID cells with *HUWE1* mutations.

**Figure 1.**
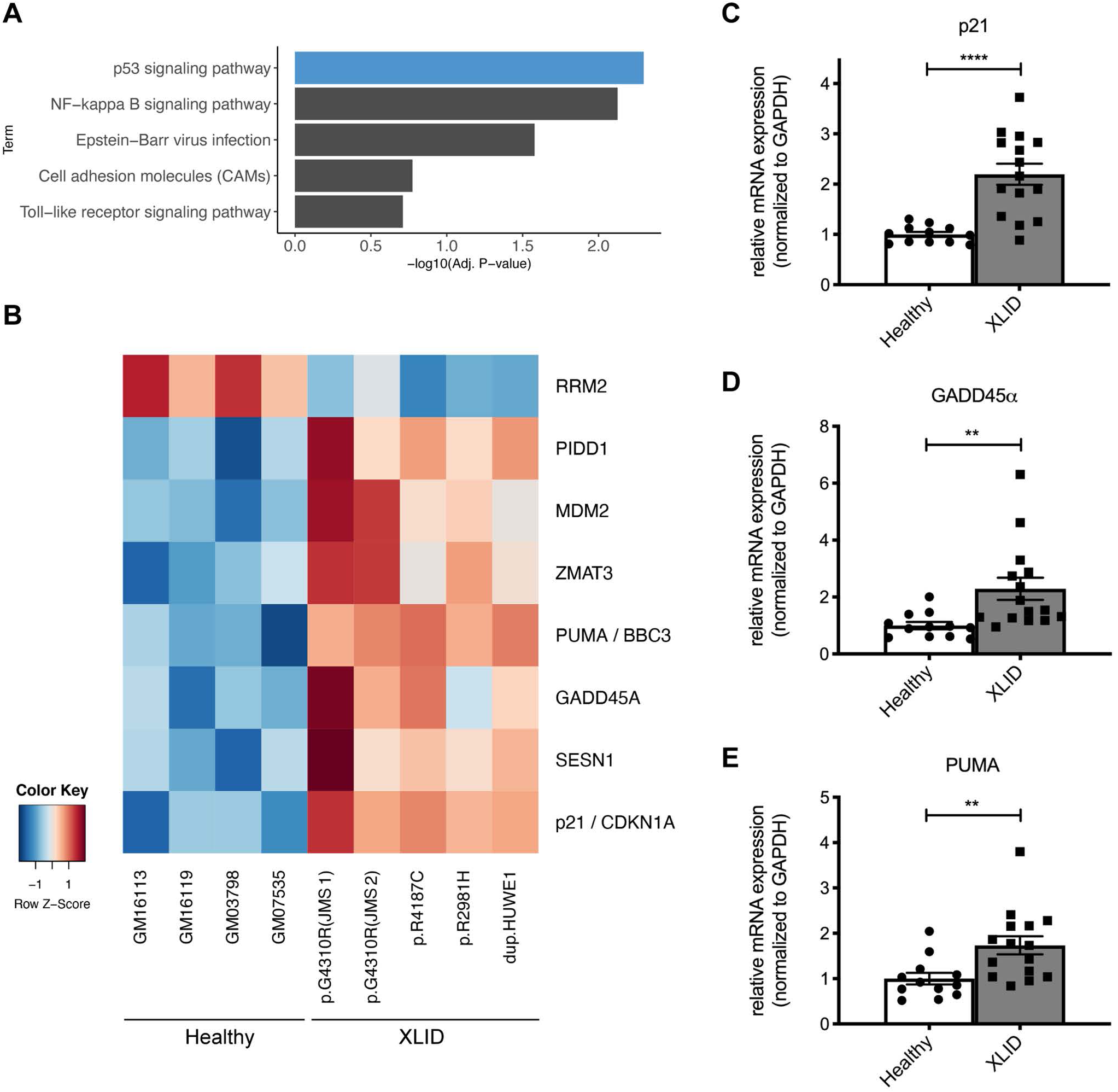
p53 signalling is hyperactivated in cells from XLID patients with mutated *HUWE1*. **(A)** Top five most significant KEGG pathway terms as determined by the Gene Set Enrichment Analysis (GSEA) of common differentially expressed genes in XLID patient-derived lymphoblastoid cells (LCLs) compared to healthy individual cells, (Benjamini corrected p-value <0.05). **(B)** Heatmap of common differentially expressed genes in XLID compared to healthy LCLs, belonging to the KEGG p53 signalling term. **(C-E)** mRNA levels of p53 target genes *p21* (C), *GADD45α* (D) and *PUMA* (E), determined by RT-qPCR analysis of four healthy (GM03798, GM07535, GM16113, GM16119) and five XLID LCLs with mutated HUWE1 (p.R2981H, p.R4187C and p.G4310R, belonging to JMS) and *HUWE1* duplication. All error bars indicate mean ± SEM (n = 3). Two-tailed unpaired t-test; **p ≤ 0.01, ****p ≤ 0.0001. See also Figure S1.

### Deregulated expression of p53 target genes contributes to altered cell cycle and reduced proliferation in HUWE1-promoted JMS

To explore the impact of increased p53 signaling (Figure 1) on cellular functioning, we focused on one of the most severe XLID forms the JMS, caused by mutated HUWE1 p.G4310R. We confirm that p53 is hyperactive in JMS, by detecting significantly increased mRNA levels of p53 targets PUMA and GADD45α in two independent JMS XLID patient-derived LCLs compared to the healthy WT control (Figure S1A and S1B). The subsequent immunoblot analysis of JMS LCLs revealed significantly increased p21 protein levels (Figure S1C), suggesting that increased p53 activity is directly reflected in deregulated protein status of the p53 targets. In line with the previous work (4), HUWE1 p.G4310R protein levels were reduced, while p53 consequently accumulated in the JMS LCLs (Figure S1C). Interestingly, on a functional level the increased p53 signaling was accompanied by impaired cell cycle progression, with significant accumulation of JMS LCLs in the G1-phase and reduction in the S-phase of the cell cycle (Figure S1D). Besides cell cycle impairments, a slight increase in the apoptosis rate measured by Annexin V staining (Figure S1E), and significantly reduced proliferation (Figure S1F) were observed in the two JMS LCLs. Taken together, these findings suggest that in differentiated JMS cells, such as LCLs, increased p53 signaling results in impaired cell cycle progression, increased apoptosis and reduced proliferation.

### Increased p53 signaling leads to failed neural differentiation of JMS hiPSCs

To determine if elevated p53 signaling observed in Figure 1. affects the JMS neurodevelopment, we next modeled the disease in a set of neural induction experiments using human induced pluripotent stem cells (hiPSCs). The JMS hiPSCs, generated by reprograming of JMS patient-derived fibroblasts (III-2)(22), were characterized by unperturbed expression of pluripotency markers SSEA4 and OCT4, and embryoid body (EB) formation comparable to the healthy individual (WT) hiPSCs (Figure S2A – S2D). Interestingly, no difference in the cell cycle progression was observed between WT and JMS hiPSCs (Figure S2E), despite increased p53 levels in JMS hiPSCs (Figure S5). To model JMS we next performed neural induction experiments up to stage of rosettes, using a modified protocol for the generation of telencephalic organoids (23). While both WT and JMS hiPSCs resulted in comparable EB formation (Figure 2A, and Figure S2D), JMS hiPSCs failed to undergo neural differentiation, forming only a very limited number of neural rosettes (Figure 2B). This observation is supported by the mRNA expression profiles, which upon neural induction revealed dramatically reduced mRNA expression levels of neural differentiation markers *DCX, TUJ1* and *MAP2* in JMS (Figure 2E and 2F, and Figure S3C). The levels of: the pluripotency markers *OCT4* and *NANOG*, the marker of proliferating NPCs *NESTIN*, and the neuroectoderm cell fate determinant *PAX6*, did not significantly differ between WT and JMS (Figure 2C and 2D, and Figure S3A and S3B). In addition, TUJ1 protein expression and localization were dramatically altered in JMS rosettes, when compared to WT samples (Figure 2H). The protein expression differences were less pronounced in the case of NESTIN (Figure 2G), consistent with mRNA expression analysis (Figure 2D). To determine the extent to which impairments in neural differentiation correlate with p53 signaling, the expression of several important p53 targets was analyzed next. Interestingly, while no difference was observed in the expression of *p21, GADD45α, PUMA* and *BAX* in the hiPSCs, levels of all tested p53 targets were significantly increased upon neural induction in JMS cells, when compared to WT cells (Figure 2I – 2L), thereby indicating elevated p53 signaling upon neural induction of JMS hiPSCs. In summary, our results suggest that capacity to undergo neural differentiation is significantly reduced in JMS and accompanied by elevated p53 signaling, while the stem cell identity is unaltered.

**Figure 2.**
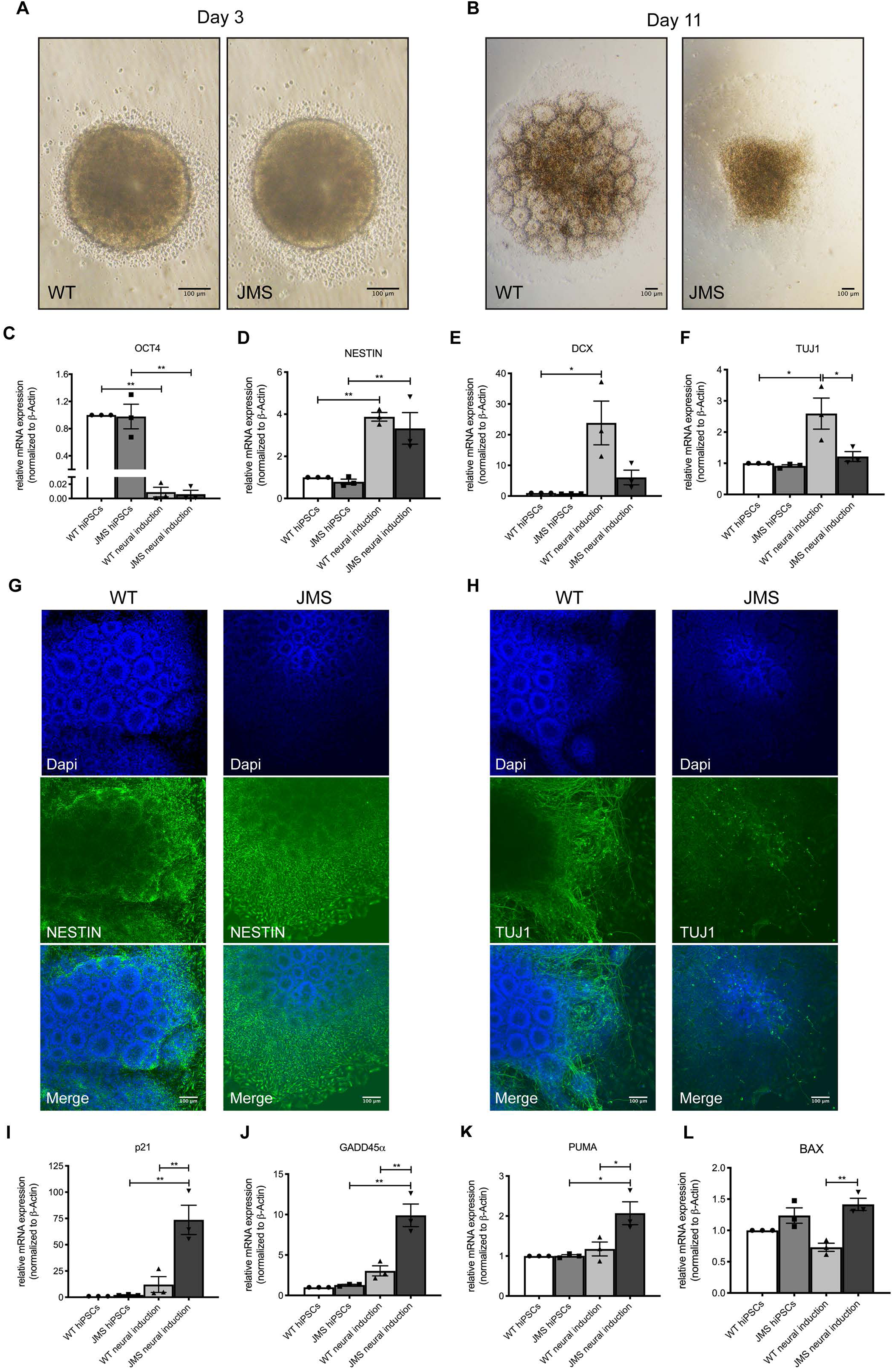
Neural differentiation of JMS hiPSCs is impaired and accompanied by activation of p53 signaling. (**A and B**) Representative bright-field images of neural differentiation of WT and JMS hiPSCs at day 3 (A) and day 11 (B); EBs and neural rosette structures are visible, respectively. (**C – F**) RT-qPCR analysis of mRNA expression of: *OCT4* (C), *NESTIN* (D), *DCX* (E) and *TUJ1* (F) genes in WT and JMS hiPSCs and neural cells (collected at day 13). (**G and H**) Immunofluorescence analysis of NESTIN (G) and TUJ1 (H) in WT and JMS hiPSCs at day 13 of neural differentiation. (**I – L**) mRNA levels of the p53 target genes p21 (I), GADD45α (J), PUMA (K) and BAX (L) in WT and JMS hiPSCs and neural cells (collected at day 13) addressed by RT-qPCR. All error bars indicate mean ± SEM (n = 3); One-way ANOVA followed by Bonferroni post-test; *p ≤ 0.05, **p ≤ 0.01, n.s ≥ 0.05. Scale bar: 100µm. See also Figure S2 and S3.

### JMS cerebral organoids fail to fully develop

To assess the extent to which observed neural differentiation impairments influence brain organogenesis, JMS cerebral organoids were developed using hiPSCs. While no differences were observed at the stage of EB formation and growth (Day 4), major developmental failure occurred in JMS cerebral organoids upon formation of neuroepithelial buds (Day 14) (Figure 3A, and Figure S4). Majority of JMS organoids failed to develop further and only very few (1-2 out of initial 96 in several independent experiments) passed Day 35 of development. The JMS organoids that did develop were significantly smaller than the WT organoids at Days 14 - 60 (Figure 3A, Figure S4). Further, the regions of cortical layering were dramatically reduced in the JMS organoids and exhibited reduced cellularity and increased ventricle-like spaces (Figure 3B). The reduced cellularity was accompanied by visibly decreased Ki67 staining, a marker of proliferation, in JMS organoids (Figure 3C and 3D). In contrast to WT organoids where majority of Ki67 signal was distributed in the proliferating region, the Ki67 signal in JMS organoids was ectopic (Figure 3C and 3D), thus corresponding to altered cortical layering observed in Figure 3B. Taken together, reduced cellularity, altered cortical layering and decreased size, closely recapitulate symptoms described in JMS patients, including microcephaly, enlarged ventricles and reduced life span (4).

**Figure 3.**
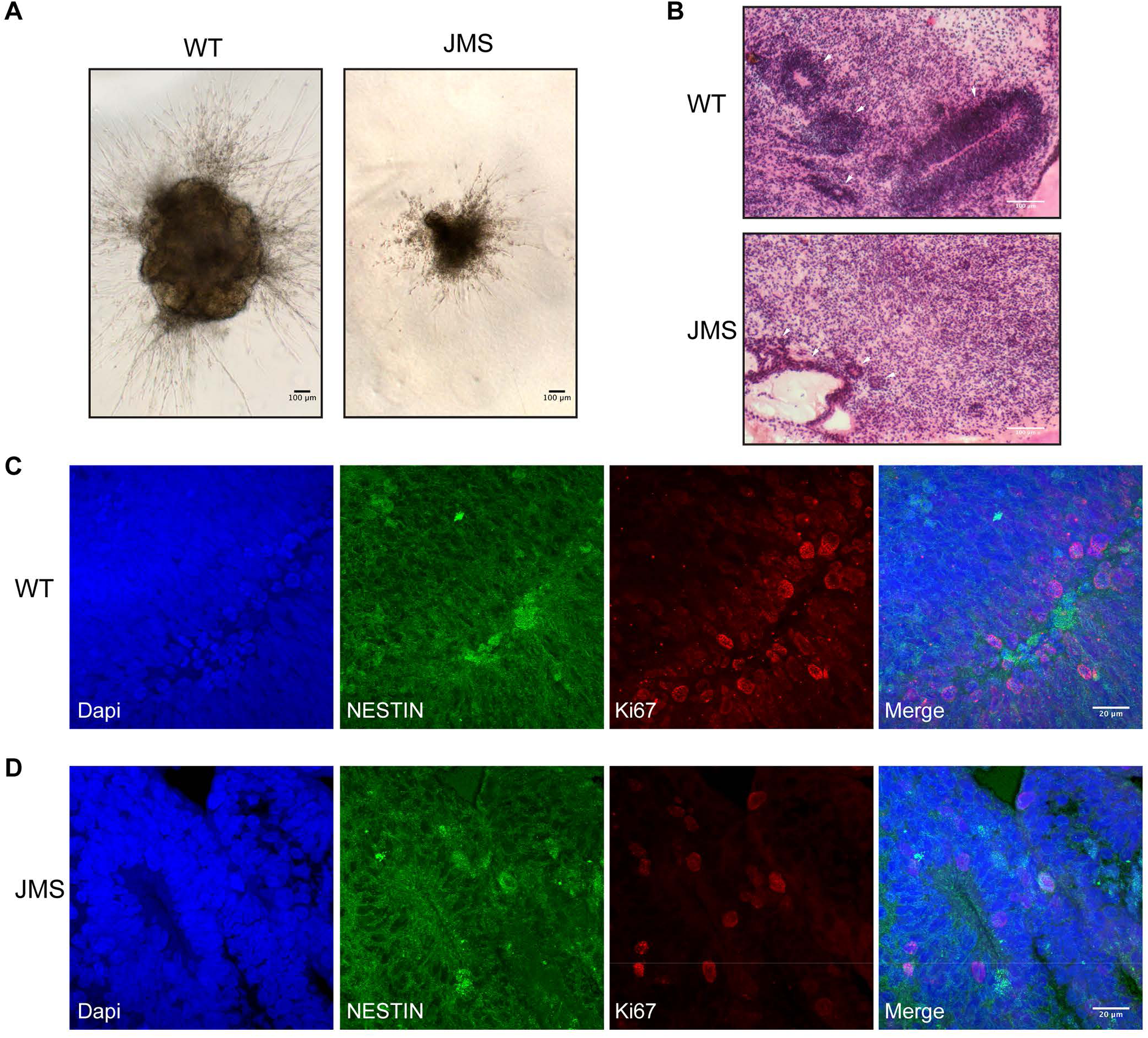
Cerebral organoids derived from JMS hiPSCs are reduced in size and exhibit altered cellular organization. (**A**) Representative bright-field images of WT and JMS cerebral organoids at day 14 of differentiation. Scale bar: 100µm. (**B**) Hematoxylin Eosin Saffron (HES) staining of 60 days old WT and JMS cerebral organoid cryosections. Scale bar: 100µm. (**C and D**) Immunofluorescence analysis of NESTIN and Ki67 in 60 days old WT (C) and JMS (D) cerebral organoids. Scale bar: 20µm. See also Figure S4.

### p53 down-regulation rescues neurodevelopmental defects in JMS

As shown in Figure 2, the impaired neural differentiation in JMS was accompanied by elevated p53 signaling. However, to which extent elevated p53 levels directly cause neurodevelopmental impairments in JMS disorder remained unclear. To address this and test if neurodevelopmental defects are a consequence of increased p53, we infected JMS hiPSCs with lentivirus encoding scrambled control shRNA (shControl), or shRNA targeting p53 (shp53), and confirmed knock-down efficiency (Figure S5). Importantly, p53 knock-down resulted in JMS neural rosette formation comparable to the WT (Figure 4A and 4D), thus efficiently rescuing impaired neural differentiation observed in JMS and JMS shControl hiPSCs (Figure 4B and 4C). Rescue of neural differentiation in JMS shp53 rosettes was accompanied by increased expression of maturation markers *TUJ1* and *DCX* in JMS shp53 rosettes, to the level significantly higher than in JMS and JMS shControl samples (Figure 4G and 4H). As expected, relative expression of *NESTIN* did not significantly differ between the four hiPSCs lines (Figure 4E). Interestingly, the expression of *PAX6*, the neuroectoderm cell fate determinant, was significantly higher in JMS shp53 rosettes when compared to JMS and JMS shControl (Figure 4F). Besides the impact on maturation markers, p53 knock-down resulted in significantly reduced expression of p53 targets *p21* and *GADD45α* (Figure 4I and 4J). Similarly, levels of *PUMA* and *BAX* (Figure 4K and 4L) were reduced, however the reduction was not significant. In summary, p53 knock-down in the JMS hiPSCs with destabilized HUWE1 p.G4310R, resulted in a striking rescue of the capacity to produce maturing neurons, thus strongly suggesting that elevated p53 signaling is a predominant cause of impaired JMS neural differentiation.

**Figure 4.**
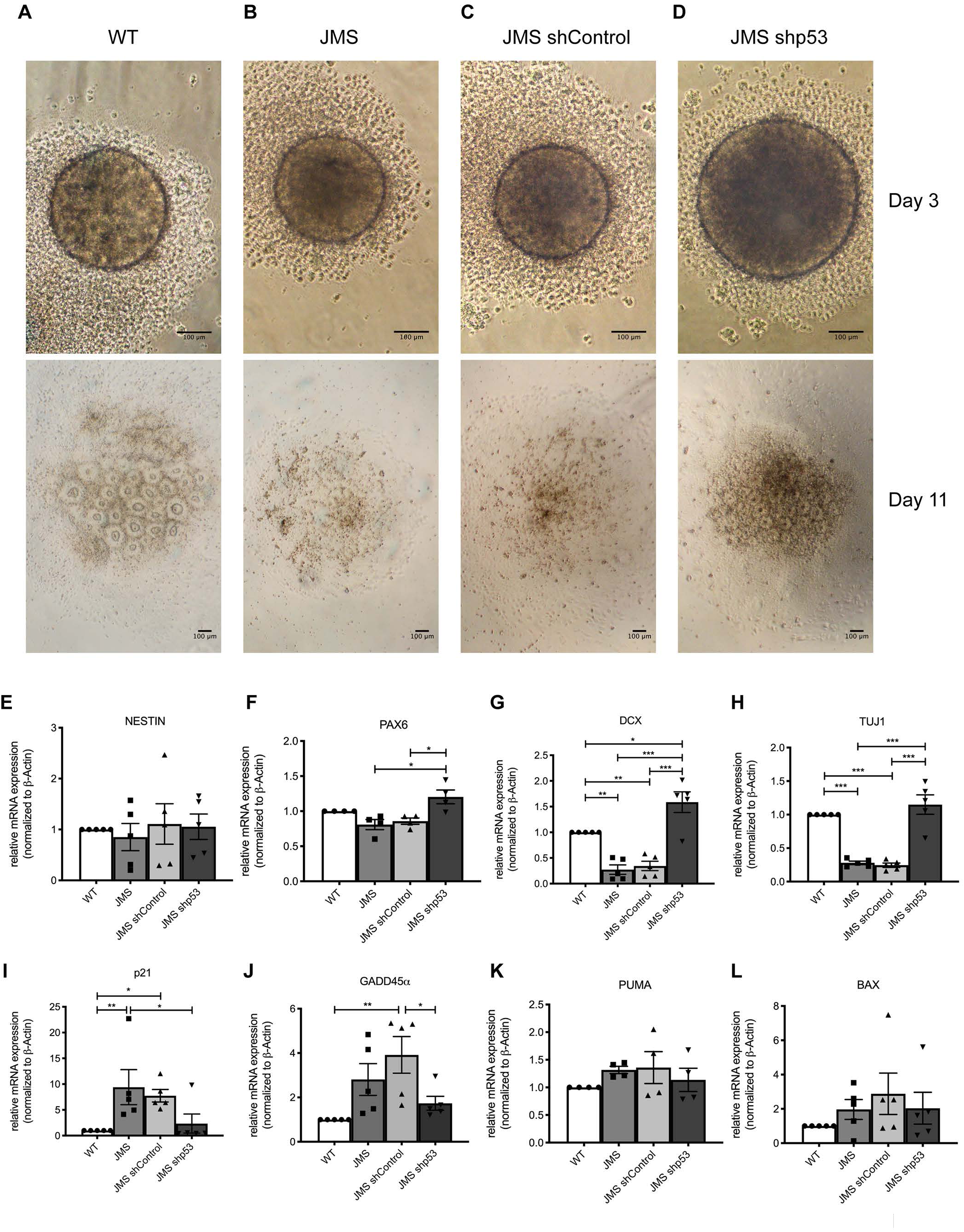
p53 down-regulation rescues neurodevelopmental defects in XLID JMS. (**A-D**) Representative bright-field images of neural differentiation of WT (A), JMS (B), JMS shControl (C) and JMS shp53 (D) hiPSC lines at day 3 and day 11. Scale bar: 100µm. (**E-L**) mRNA expression levels of *NESTIN* (E), *PAX6* (F), *DCX* (G), *TUJ1* (H), *p21* (I), *GADD45α* (J), *PUMA* (K) and *BAX* (L) genes in WT and JMS neural cells (collected at day 13) addressed by RT-qPCR. All error bars indicate mean ± SEM (n ≥ 3); one-way ANOVA followed by Bonferroni post-test; *p ≤ 0.05, **p ≤ 0.01, ***p ≤ 0.001, ****p ≤ 0.0001, n.s ≥ 0.05. See also Figure S5.

## DISCUSSION

HUWE1 is an essential E3 ubiquitin ligase that plays a decisive role in early stages of neurodevelopment (7, 24). In humans *HUWE1* mutations associate with XLIDs, however the underlying pathomechanisms are still poorly understood. In this work we demonstrate that different *HUWE1* mutations result in hyperactivation of p53 signaling, which causes impaired neural differentiation in XLID. Specifically, RNA sequencing performed on cells from five independent XLID patients with different *HUWE1* mutations, identified p53 signaling as the most significantly altered pathway, when compared to four healthy control cell lines (Figure 1). Additional analysis of individual genes revealed that p53 signaling is increased in the XLID cells. These findings support the hypothesis that p53 activation might be a unique link between different genetic developmental disorders (13). Having established that the common feature of HUWE1-related XLIDs is deregulation of p53 pathway, we next focused on JMS, one of the most severe forms of XLID. In addition to increased p53 signaling, JMS is characterized by reduced HUWE1 p.G4310R levels and p53 accumulation (4) (Figure S1). By recapitulating the XLID JMS and healthy individual neurodevelopment, we show that increased p53 signaling significantly reduces the JMS hiPSCs capacity to undergo neural differentiation, while the ability to form EBs and produce NPCs remains unaffected (Figure 2). Interestingly, recent work focusing on an XLID-causing mutation in another E3 ubiquitin ligase RNF12 indicated that, similarly to HUWE1 p.G4310R, loss of RNF12 function results in altered neural differentiation (25). The reduced neural differentiation capacity observed in HUWE1-promoted JMS (Figure 2) is further supported by studies in mice, which indicated the crucial importance of Huwe1 in cell cycle exit of NPCs and their subsequent neurogenic differentiation in both cerebral cortex (9) and cerebellum (10). Interestingly, the impaired JMS neurogenic capacity was accompanied by increased levels of the p53 target genes: *p21, GADD45α, PUMA* and *BAX* at the final stages of neural differentiation (Figure 2I – 2L), similar as in LCLs (Figure S1), thus indicating that p53 transcriptional activity is elevated in differentiated JMS cells. Besides p53 protein accumulation, specific patterns of posttranslational modifications, such as phosphorylation and acetylation, promote p53 signaling and the expression of p53 target genes (26). While we did not detect significant increase in p53 Serine 15 phosphorlytion in JMS cells (data not presented), it will be interesting in the follow-up work to determine modifications and sites on p53 at which they occur, to further explore p53 activity and regulation during JMS neurodevelopment. The impaired neural differentiation, resulting from p53 accumulation in JMS, was reflected in severely perturbed brain organogenesis as demonstrated by development of JMS cerebral organoids (Figure 3). Only very few of the JMS organoids that did develop had decreased size, which is in line with the microcephaly observed in JMS patients (4), and were characterized by reduced cellularity and lack of the laminar patterning, features reported previously in the mouse Huwe1 mouse models (9, 10). Interestingly, the observation that elevated p53 activity accompanies neurodevelopmental impairments in JMS is supported by recent studies showing that patients with germline *TP5*3 mutations present with microcephaly (27), as well as that microcephaly and neurodevelopmental defects in several human neurodevelopmental disorders are p53-dependent (13). This supports the idea that deregulation of p53 activity could have central role in the onset of various neurological conditions. Notably, p53 knock-down and consequently restored p53 signaling in the JMS hiPSCs resulted in a striking rescue of the capacity to produce maturing neurons (Figure 4), thus strongly suggesting that impaired neural differentiation in JMS is predominantly caused by p53 hyperactivation. Remarkably, recent work showed that depletion of Huwe1 in *Drosophila* results in impaired development, characterized by aberrant cell-cycle phasing and failure to enter S-phase; similarly to our observations, all the Huwe1 loss-of-function phenotypes were successfully suppressed by p53 knockdown (28). In mouse iPSCs p53 accumulation was furthermore shown to cause impaired neural differentiation, which was successfully overcome by p53 inactivation (29). In summary, our findings suggest that p53 deregulation is the common process underlying wide range of *HUWE1*-causing XLID syndromes. While p53 targeting during development is problematic, identification of specific p53 targets which could rescue developmental phenotypes could open avenues to therapies in XLID and possibly other neurodevelopmental disorders. Accordingly, since different HUWE1-promoted XLID syndromes present with unique symptoms, subsequent studies are needed to clarify how increased p53 signaling contributes to different phenotypes.

Taken together, the findings presented in this study demonstrate that HUWE1 plays vital role in regulation of p53 signaling during human neurodevelopment and that HUWE1-p53 pathway is crucial for enabling NPCs to undergo terminal differentiation, an imbalance of which causes the onset of the neurodevelopmental disorder XLID.

## MATERIALS AND METHODS

### Cell culture

The healthy individual LCLs (GM03798, GM07535, GM16113, GM16119) were obtained from Coriell Cell Repository (Coriell Institute for Medical Research, USA). The XLID LCLs with HUWE1 duplication, or HUWE1 p.R2981H and p.R4187C mutations were published previously (3). The JMS1 and JMS2 LCLs with mutation HUWE1 p.G4310R were obtained from the individuals III-2 and IV-4, and WT LCLs from healthy relative IV-1, from the original family reported by Juberg and Marsidi (4, 22). Informed consent was obtained from each participant enrolled in the study of X-linked Intellectual Disability according to the regulations of the Institutional Review Board of Self Regional Healthcare and Greenwood Genetic Centre, South Carolina, USA. All LCLs were maintained in RPMI 1640 (Corning™) with 15% FBS (ThermoFisher Scientific), 1% penicillin/streptomycin (Sigma). Patient-derived human induced pluripotent stem cells (hiPSCs) were generated by Applied StemCell (California, USA) by viral-free episomal reprogramming of JMS individual III-2 fibroblasts trough electroporation with three reprogramming plasmids containing human sequences for *OCT4, SOX2, KLF4, LIN28, L-MYC*. The obtained hiPSC clones were grown in mTeSR1 medium (STEMCELL Technologies), on CellMatrix Basement Membrane Gel (ATCC) coated dishes, as well as the healthy individual hiPSCs (ATCC-DYS0100, ATCC, Manassas, USA). JMS hiPSCs were characterized by alkaline phosphatase staining (data not shown) and immunofluorescence analysis of pluripotency markers OCT4 and SSEA4 (Figure S2) and gene expression analysis (*OCT*4, *SOX2, NANOG*) (data not shown). All cell lines were grown in a humidified 5% CO_2_ atmosphere at 37°C, and regularly tested for mycoplasma.

### RNA isolation and quantitative PCR

RNA isolation was performed using RNeasy mini kit (Qiagen) and cDNA subsequently generated with MultiScribe™ Reverse Transcriptase (ThermoFisher Scientific) according to the manufacturer’s instructions. Quantitative PCR (qPCR) was performed with Power SYBR™ Green PCR Master Mix (Applied Biosystems™) on a StepOnePlus™ Real-Time PCR System. qPCR was performed with primer pairs (Sigma) listed in Table S1. The relative expression levels were determined by normalization to *GAPDH* or *β-Actin*, as indicated in the figure legends.

### RNA sequencing

RNA sequencing libraries were generated using SENSE mRNA-Seq library prep kit according to manufacturer’s instructions (Lexogen GmbH, Vienna, Austria). In brief, 300 ng of total RNA was prepared and incubated with magnetic beads coated with oligo-dT, then all other RNAs except mRNA were removed by washing. Library preparation was then initiated by random hybridization of starter/stopper heterodimers to the poly(A) RNA still bound to the magnetic beads. These starter/stopper heterodimers contain Illumina-compatible linker sequences. A single-tube reverse transcription and ligation reaction extends the starter to the next hybridized heterodimer, where the newly-synthesized cDNA insert was ligated to the stopper. Second-strand synthesis was performed to release the library from the beads. The resulting double-stranded library was purified and amplified (13 PCR cycles) prior to adding the adaptors and indexes. Finally, libraries were purified using the Mag-Bind RXNPure Plus beads (Omega Bio-tec, GA, USA), quantitated by qPCR using KAPA Library Quantification Kit (Kapa Biosystems, Inc., MA, USA) and validated using Agilent High Sensitivity DNA Kit on a Bioanalyzer (Agilent Technologies, CA, USA). The size range of the DNA fragments were measured to be in the range of app. 200-450 bp and peaked around 270 bp.

Libraries were normalized and pooled to 2.4 nM and subject to clustering (by a cBot Cluster Generation System on HiSeq4000 flowcells (Illumina Inc., CA, USA), according to manufacturer’s instructions. The sequencing (75 cycles single end reads) were performed on an Illumina HiSeq4000 instrument, in accordance with the manufacturer’s instructions (Illumina, Inc., CA, USA). FASTQ files were created with bcl2fastq 2.20.0.422 (Illumina, Inc., CA, USA).

### Bioinformatic analysis

#### RNA-seq analysis

Raw reads were demultiplexed and mapped to the human genome (Ensembl GRCh38.84) using STAR aligner v2.4.0 (30). Read counts per gene were calculated with htseq-count (31) v0.6.0, with features being counted at the exon level. Normalization and differential expression was analyzed with limma voom (32). Read counts were filtered to remove genes with less than an average of 1 read per sample post-normalization. Fold changes >1 and a FDR (false discovery rate) <0.3 were considered significant.

#### Functional Annotation

Pathway analysis of differentially expressed genes (DEGs) with fold change >1 and FDR <0.3 was performed using the functional annotation tool DAVID (Database for Annotation, Visualization, and Integrated Discovery, v6.8) (33). Significance of KEGG_PATHWAY terms was evaluated by hypergeometric testing (as a function in DAVID), using p-values with Benjamini correction for multiple hypothesis testing. Pathways were considered significantly enriched with a Benjamini corrected p-value <0.05.

#### Heatmap

Log-CPM (copies per million) normalized expression values of DEGs that appeared on the KEGG p53 signalling pathway list were used as input to the R package ‘pheatmap’ v1.0.12 (R version 3.4.1) to create a heatmap. Log-CPM values were scaled across rows (genes) to generate Z-scores, which were then used for colouring the heatmap.

### Whole cell extracts

LCLs were collected and washed twice with ice cold PBS, followed by flash freezing in liquid N2. Whole cell extracts (WCEs) were obtained similarly as described previously (34). Briefly, cell pellets were resuspended in lysis buffer I (10 mM Tris-HCl pH 7.8, 200 mM KCl, 0.1 mM MG-132, 1 μM PMSF, halt protease and phosphatase inhibitor cocktail (Thermo Scientific)); followed by addition of lysis buffer II (10 mM Tris-HCl pH 7.8, 600 mM KCl, 2 mM EDTA, 40% glycerol, 0.2% NP-40, 0.1 mM MG-132, 1 μM PMSF, halt protease and phosphatase inhibitor cocktail); rotated 30 minutes at 4 °C, sonicated and centrifuged. The supernatants representing WCEs were collected. hiPSCs WCEs were prepared as described in (35). Briefly, hiPSCs were resuspended in two packed cell volumes (PCV) hypotonic lysis buffer (20 mM HEPES at pH 7.9, 2 mM MgCl2, 0.2 mM EGTA, 10% (v/v) glycerol, 0.1 mM PMSF, 2 mM DTT, complemented with Halt Protease and Phosphatase Inhibitors Cocktail) and incubated 5 min at 4 °C, followed by three freeze/thaw cycles. NaCl and Nonidet-P40 were added to final concentration 0.5 M and 0.5% (v/v), respectively, and samples incubated 20 min at 4 °C, sonicated and centrifuged. The supernatants were diluted with eight PCV of lysis buffer containing 50 mM NaCl and used for subsequent analysis.

### Immunoblot analysis

WCE proteins were separated on 4–12% Bis–Tris polyacrylamide gels (Invitrogen) followed by transfer to Amersham^™^ Hybond® P Western blotting membranes, PVDF (Sigma). Primary antibodies HUWE1 (A300-486A, Bethyl Laboratories), p53 (MA5-12571, ThermoFisher Scientific), p21 (33-7000, ThermoFisher Scientific), Tubulin (T9026, Sigma-Aldrich), β-Actin (A1978, Sigma-Aldrich)) were detected using infrared (IR) Dye-conjugated secondary antibodies (Li-COR Bioscienecs) and the signal visualized by Odyssey Scanner, LI-COR Biosciences.

### Cell cycle analysis

Cells were fixed with 70% ethanol, stained with propidium iodide (Sigma Aldrich) at a final concentration of 20 µg/ml in PBS with the addition of 100 µg/ml RNase A (37°C, 30 min in the dark). Flow cytometry analysis was performed with the BD FACS CANTO SYSTEM (BD Biosciences). Data were analyzed in FlowJo LLC software (USA).

### Analysis of apoptosis

The apoptotic WT, JMS1 and JMS2 LCLs were identified using Dead Cell Apoptosis Kit (Molecular Probes). WT LCLs treated with 250 μM H_2_O_2_, for 2h, at 37 °C served as a positive control. LCLs were washed with cold PBS and diluted to 1 × 10^6^ cells/mL in 1X annexin-binding buffer; 100 μl of cell suspension was labeled with Annexin V Alexa Fluor^®^ 488 and Propidium Iodide (Sigma), incubated for 15 minutes at room temperature and analyzed on a BD FACSAria II (BD Biosciences). The apoptotic cell fraction was determined by FlowJo, LLC software (USA).

### Cell proliferation

LCLs (1.5 x 10^4^ per well) were seeded in 96 well plates and growth rate monitored by counting every 22h (Countess II Automated cell counter, ThermoFisher Scientific).

### Neural differentiation

Neural differentiation was performed according to modified protocol (23). WT and JMS hiPSC colonies were pre-incubated with 10 μM ROCK inhibitor (Y-27632, Millipore) for 1h, and dissociated with Accutase (STEMCELL Technologies). Cells were re-aggregated (10000 cells per well) in U-bottom ultralow attachment 96-well plates (Corning) and cultured in “EB-medium” (DMEM/F12-GLUTAMAX medium, 2% B27 supplement without vitamin A (Invitrogen), 1% N2 supplement (Invitrogen), 50 μM 2-mercaptoethanol (2-ME), 5μM Y-27632 and 100 ng/ml recombinant mouse Noggin (R&D Systems, 1967-NG-025)). After 2 days half of the culture medium was replaced with “EB-medium” with vitamin A. After 2 additional days free-floating EBs were transferred on Matrigel (BD Biosciences) coated plates and cultured in neural medium (DMEM/F12-GlutaMAX containing 2% B27 with Vitamine A, 1% N2, 50 uM 2-mercaptoethanol) for 2 days. Terminal differentiation was induced using a NEUROBASAL-A medium (ThermoFisher Scientific) with 1% N2, 2% B27 with vitamin A, 1% Glutamax, 1% nonessential amino acids (NEAA) and 50 μM 2-Mercaptoethanol, 200 nM ascorbic acid, 10 ng/ml BDNF (R&D), 10 ng/ml GDNF (R&D) and 1 mM dibutyryl-cAMP (Sigma), and medium exchanged every 48h. After 6 days of terminal differentiation the neural cells were collected for subsequent immunostaning or qPCR analysis.

### shRNA knockdown

WT and JMS hiPSCs were transduced with lentiviruses encloding for the non-specific control short-hairpin RNA (shControl; pLKO.1 puro (Addgene ID: 8453)) or the p53 targeting shRNA (shp53 pLKO.1 puro shRNA (Addgene ID: 19119) (36). 72 hours after infection, puromycin selection (1 µg/ml) was started and carried out for 48 hours. Stable transduced colonies were expanded and p53 down-regulation assessed by immunoblot analysis.

### Cerebral organoids

hiPSCs have been differentiated to form cerebellar organoids following protocol described in (37). Briefly, WT and JMS hiPSCs were dissociated and induced to form EBs, which after six days were transferred in neural induction medium allowing neuroectoderm formation. Neural ectoderm induced EBs were embedded in Matrigel (BD Biosciences, 356234) and, upon outgrowth of neuroepithelaial buds, transferred to spinning bioreactor. 60 days after the initiation of differentiation cerebral organoids were collected and prepared for cryosectioning.

### Immunostaining

WT and JMS neural cells at day 13 (Figure 2G and 2H), or hiPSCs WT, JMS-clone a and JMS-clone b (Figure S2) were fixed for 15 minutes with 4% paraformaldehyde or ice cold methanol, respectively. Upon permeabilized at RT with 0.1% Triton-X in 1X PBS for 15 min, samples were incubated with 5% BSA, 5% goat gut serum, 0.1% Triton-X in 1X PBS for 45 min. Primary antibodies (OCT4 (diluted 1:200, Cell signaling C30A3); SSEA4 (diluted 1:200, Cell signalling MC813), NESTIN (diluted 1:400, Abcam ab2203), TUJ1 (diluted 1:750, Covance MMS-435P), Ki67 (diluted 1:200, Abcam ab15580) diluted in 0.5% BSA, 0.5% goat gut serum, 0.1% Tween-20 in 1X PBS (PBS-T), were added to samples and incubated over-night at 4 °C. After washing with PBS-T, samples were incubated with fluorophore-conjugated secondary antibody from Invitrogen (Donkey anti-Mouse, Alexa Fluor 488, A21202; Goat anti-Rabbit, Alexa Fluor 594, Invitrogen, A11037) diluted 1:400 in 1X PBS at RT for 1 h. Upon washing with PBS-T, cells were stained with DAPI for 10 minutes at RT, and washed 3 times with PBS. The 18-µm thick cerebral organoid cryosections were immunostained following previously described procedure (11) and primary and secondary antibodies indicated above. Images of WT and JMS neural cells at day 13 of differentiation and of cerebral organoids were captured with the Zeiss LSM 510 Meta live Confocal system, using ZEN 2009 software, UV laser (405 nm) and Argon laser (488 nm). The images of neural induction experiments and cerebral organoids are a compilation of confocal Z-stacks comprising of up to 87 optical spices (3 μm intervals) and of up to 33 optical spices (0.31 um intervals), respectively into 2D using maximum intensity projection. Images of WT, JMS-clone a and JMS-clone b hiPSCs (Figure S2) were captured with the EVOS FL Auto Cell Imaging System using 4x magnification.

### Statistical analysis

To determine significance of observed changes two-tails unpaired Student t-test or ANOVA with Bonferroni post-test were used in Prism 5, as indicated in the figure legends. p< 0.05 was considered to be significant.

### Data availability

The data that support the findings of present study are available from the corresponding author on reasonable request. The RNA sequencing data reported in this paper are available in GEO under accession GSE130551.

## Acknowledgments

This work was supported by the Central Norway Regional Health Authority to B.v.L. (90270800); the Liaison Committee between the Central Norway Regional Health Authority and the Norwegian University of Science and Technology to B.v.L. (46055600-91). Supported, in part, by an Onsager Fellowship to B.v.L.; a NINDS grant (1R01NS07385A) to CES and a grant from the South Carolina Department of Disabilities and Special Needs (SCDDSN). The RNA library preparation and sequencing were performed at the Genomics Core Facility (GCF) and Microscopy was performed at the Cellular and Molecular Imaging Core Facility (CMIC), Norwegian University of Science and Technology (NTNU). GCF and CMIC are funded by the Faculty of Medicine and Health Sciences at NTNU and Central Norway Regional Health Authority. We would like to thank Ulrich Hübscher and Giovanni Maga for the inputs and discussions in the initial phase of the project.

## Author contribution

B.v.L. and C.E.S. had the original idea, and B.v.L. supervised the project. R.A. performed majority of the experiments; S.B. contributed to the RNA sequencing, and together with B.M. and D.B. to the qPCR and immunoblot analysis. B.M. and W.W. contributed to the neural induction and the immunofluorescence analysis., S.L.F.M. to the RNA sequencing analysis; E.A., Y.P. to the protein analysis, and N-B.L to FACS analysis. M.Bo., N.P.M. and C.S. contributed reagents. R.A., B.v.L, W.W. and M.Bj. interpreted the experiments. B.v.L., R.A. and B.M. wrote the manuscript and generated figures.

