## Supplemental Information for "Increased p53 signaling impairs neural differentiation causing HUWE1-promoted intellectual disabilities"

[#]joint second authors.

**Corresponding author:** *Barbara van Loon, Department of Cancer Research and Molecular Medicine, Norwegian University of Science and Technology (NTNU), Erling Skjalgssons gt 1, Trondheim 7491, Norway;; telephone +47 72 82 46 84.

### SUPPLEMENTAL TABLES AND FIGURES

#### Table S1. Primer sequences (indicated in 5'-3' direction) used for gene expression analysis by qPCR

| Target gene name | Fw/Rev | Sequence |
|---|---|---|
| *GADD45α* | Fw | GCAGGATCCTTCCATTGAGA |
| | Rev | AGCTCCTGCTCTTGGAGACC |
| *PUMA* | Fw | GTAAGGGCAGGAGTCCCAT |
| | Rev | GACGACCTCAACGCACAGTA |
| *OCT4* | Fw | TCTTTCCACCAGGCCCCCGGCTC |
| | Rev | TGCGGGCGGACATGGGGAGATCC |
| *NESTIN* | Fw | GGCGCACCTCAAGATGTCC |
| | Rev | CTTGGGGTCCTGAAAGCTG |
| *PAX6* | Fw | GCCCTCACAAACACCTACAG |
| | Rev | TCATAACTCCGCCCATTCAC |
| *TUJ1* | Fw | GCAACTACGTGGGCGACT |
| | Rev | TCGAGGCACGTACTTGTGAG |
| *DCX* | Fw | TCAGGGAGTGCGTTACATTTAC |
| | Rev | GTTGGGATTGACATTCTTGGTG |
| *BAX* | Fw | CATGTTTTCTGACGGCAACTTC |
| | Rev | AGGGCCTTGAGCACCAGTTT |
| *p21* | Fw | GGCACTCAGAGGAGGCGCCAT |
| | Rev | TAGCGCATCACAGTCGCGGC |
| *MAP2* | Fw | CAGGAGACAGAGATGAGAATTCC |
| | Rev | CAGGAGTGATGGCAGTAGAC |
| *NANOG* | Fw | AGGGTCTGCTACTGAGATGCTCTG |
| | Rev | CAACCACTGGTTTTTCTGCCACCG |
| *GAPDH* | Fw | GAGTCAACGGATTTGGTCGT |
| | Rev | TTGATTTTGGAGGGATCTCG |
| *β-actin* | Fw | AAACTGGAACGGTGAAGGTG |
| | Rev | AGAGAAGTGGGGTGGCTTTT |

**Figure S1. p53 activation is affecting cell cycle, proliferation and apoptosis in differentiated JMS patient-derived cells.** (**A and B**) mRNA levels of p53 target genes *GADD45α* (A) and *PUMA* (B) in WT, JMS1 and JMS2 LCLs, addressed by RT-qPCR. (**C**) Immunoblot analysis of the HUWE1, p53 and p21 protein levels in WT, JM1 and JM2 LCLs. Tubulin serves as loading control. (**D**) Cell cycle distribution determined by flow cytometry of WT, JM1 and JM2 LCLs. (**E**) Fraction of Annexin V-positive apoptotic cells measured by flow cytometry. (**F**) Proliferation rate of WT, JM1 and JM2 LCLs. All error bars indicate mean ± SEM ($n \geq 3$). Statistical significance determined by two-tailed unpaired t-test (A), (B), (E) and two-way ANOVA with Bonferroni post-test (C), (F); $^{*}p \leq 0.05$, $^{**}p \leq 0.01$, $^{***}p \leq 0.001$, $^{****}p \leq 0.0001$, n.s ≥ 0.05.

**Figure S2. Stemness markers and cell cycle progression are unaltered in JMS hiPSCs.** (**A-C**) Expression of the pluripotency markers SSEA and OCT4 in (A) WT, (B) JMS-clone a and (C) JMS-clone b hiPSC lines by immunofluorescence. (**D**) Representative bright-field images of WT, JMS-clone a and JMS-clone b EBs at day 3 after EB initiation (n=3). Scale bar:100µm. (**E**) Cell cycle distribution determined by flow cytometry of WT, JM-clone a and JM-clone b hiPSCs (n=3). Error bars indicate mean ± SEM.

**Figure S3. Expression of mRNA upon neural differentiation of WT and JMS hiPSCs.** (**A-C**) RT-qPCR analysis of mRNA expression levels of NANOG (A), PAX6 (B) and MAP2 (C) genes in WT and JMS hiPSCs and neural cells (collected at day 13) ($n = 3$). Error bars indicate mean ± SEM; statistical significance was calculated using the one-way ANOVA followed by Bonferroni post-test ($^{**}p < 0.01$, $^{**}p < 0.001$).

**Figure S4. Cerebral organoids derived from JMS hiPSCs have reduced size.** (**A and B**) Representative bright-field images of (A) WT and (B) JMS cerebral organoids at day 4, day 10 (when neuroectoderm is expected to be formed) and day 35 of differentiation. Scale bar:100μm.

**Figure S5. p53 knock-down in hiPSCs.** Immunoblot analysis of HUWE1 and p53 levels in WT, JMS, JMS expressing shControl and JMS expressing shp53 hiPSCs. β-actin serves as loading control.

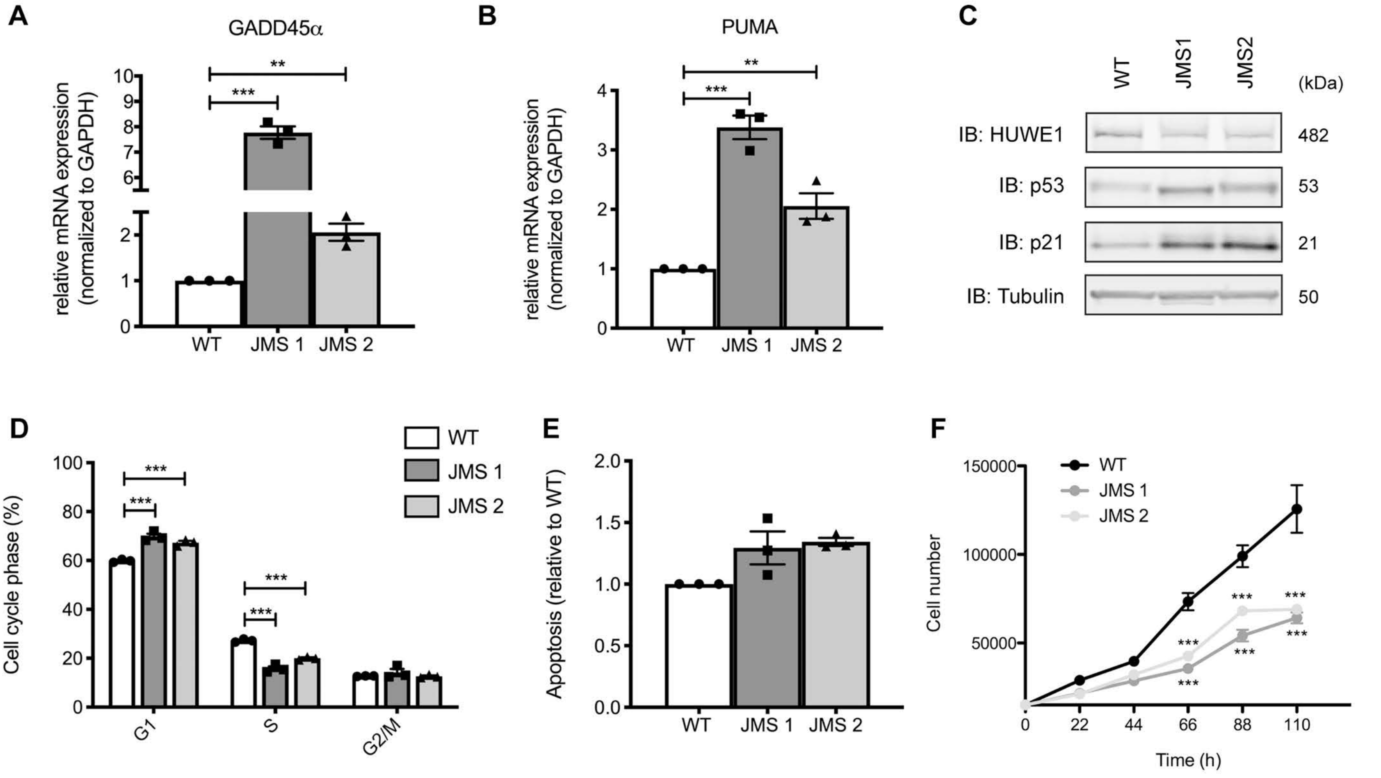
A
GADD45α
relative mRNA expression (normalized to GAPDH)
WT
JMS 1
JMS 2
B
PUMA
C
WT
JMS1
JMS2
(kDa)
IB: HUWE1
482
IB: p53
53
IB: p21
21
IB: Tubulin
50
D
Cell cycle phase (%)
G1
S
G2/M
E
Apoptosis (relative to WT)
F
Cell number
Time (h)

**Figure S1**

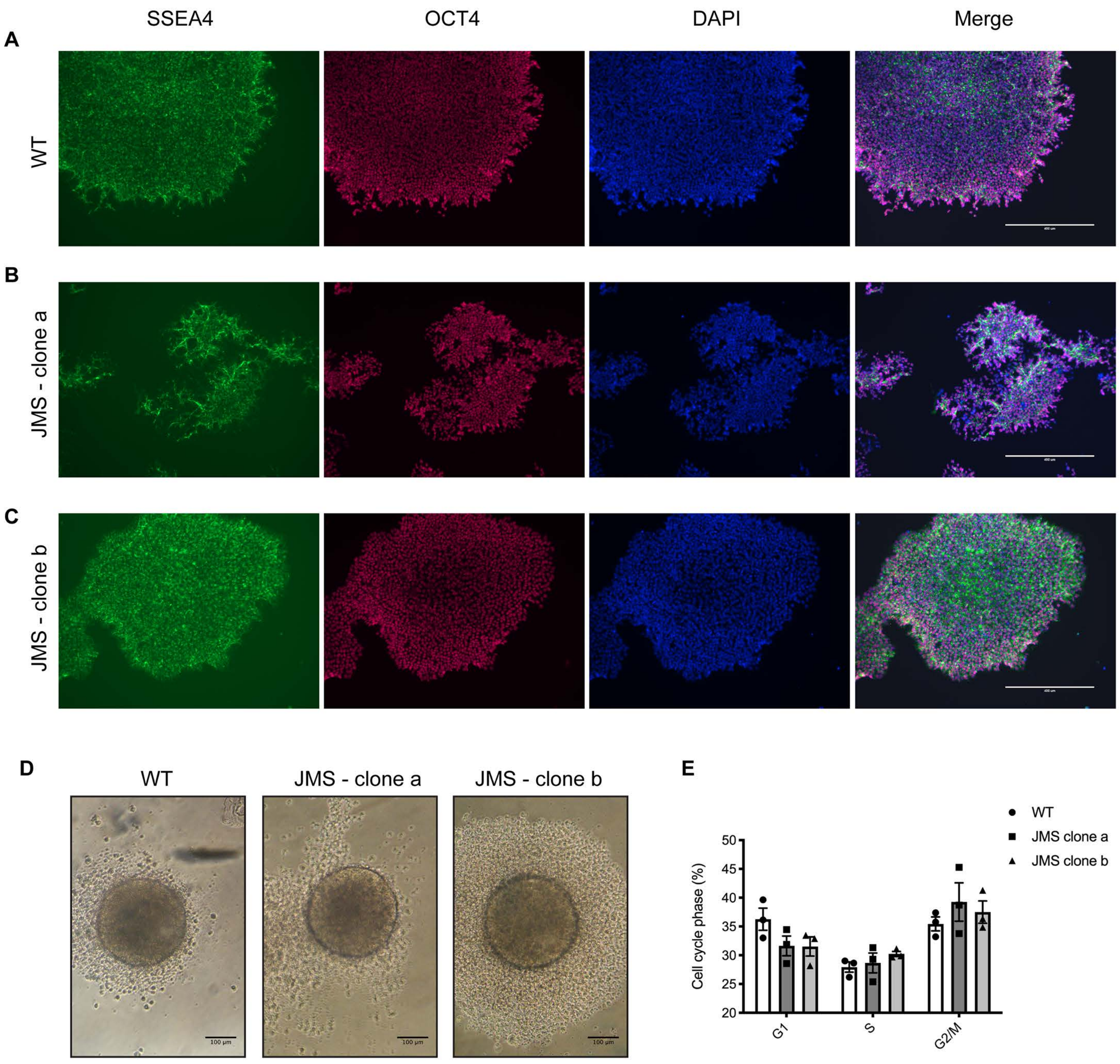

SSEA4
OCT4
DAPI
Merge
A
WT
B
JMS - clone a
C
JMS - clone b
D
WT
JMS - clone a
JMS - clone b
E
WT
JMS clone a
JMS clone b
Cell cycle phase (%)
50
45
40
35
30
25
20
G1
S
G2/M


**Figure S2**

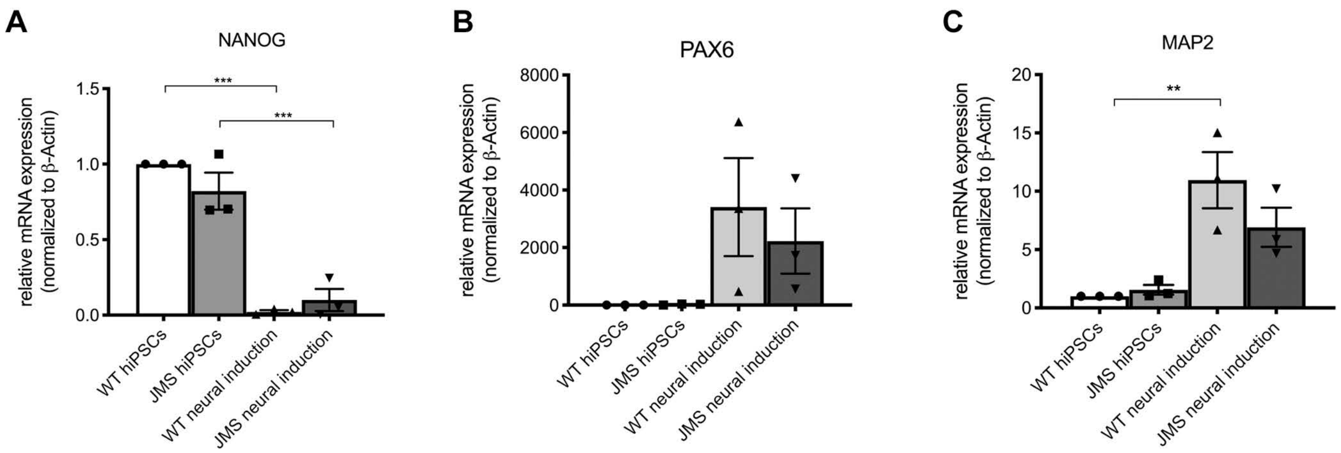
A
NANOG
relative mRNA expression
(normalized to β-Actin)
1.5
1.0
0.5
0.0
***
***
WT hiPSCs
JMS hiPSCs
WT neural induction
JMS neural induction
B
PAX6
relative mRNA expression
(normalized to β-Actin)
8000
6000
4000
2000
0
WT hiPSCs
JMS hiPSCs
WT neural induction
JMS neural induction
C
MAP2
relative mRNA expression
(normalized to β-Actin)
20
15
10
5
0
**
WT hiPSCs
JMS hiPSCs
WT neural induction
JMS neural induction

**Figure S3**

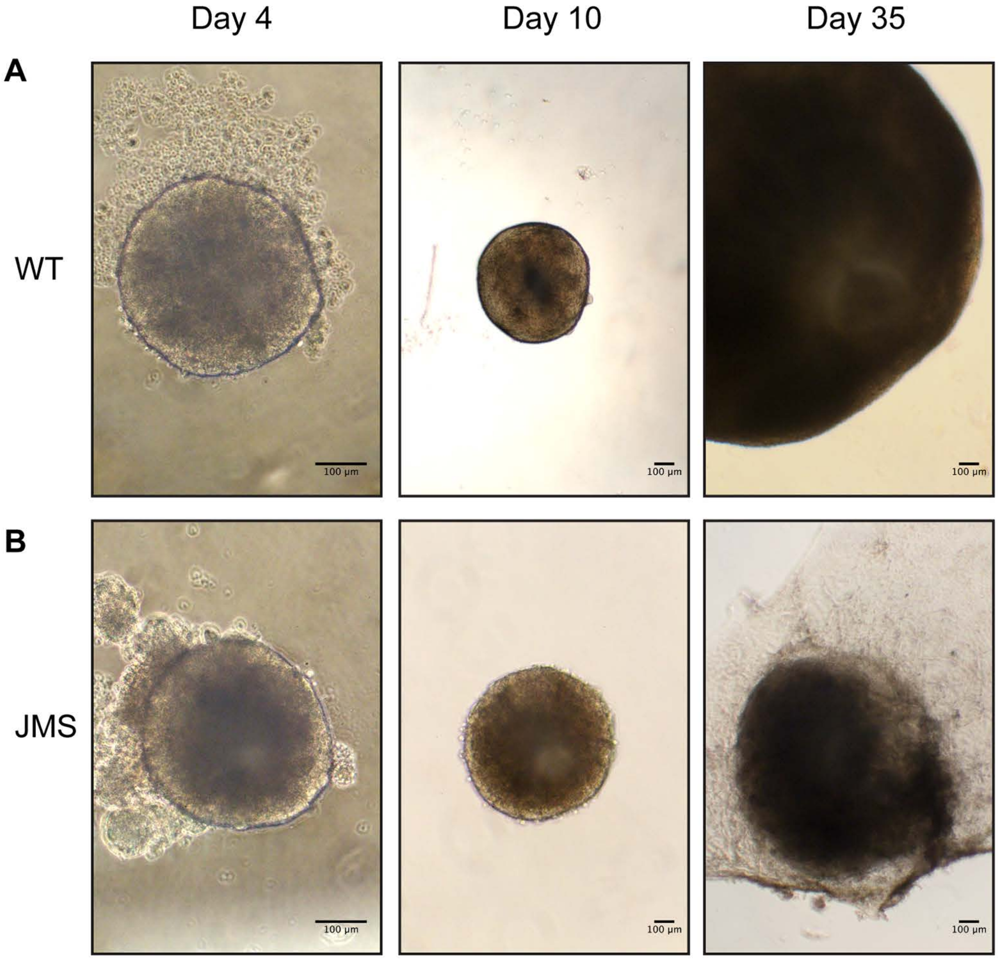
Day 4
Day 10
Day 35
A
WT
B
JMS
100 μm
100 μm
100 μm
100 μm
100 μm
100 μm

**Figure S4**

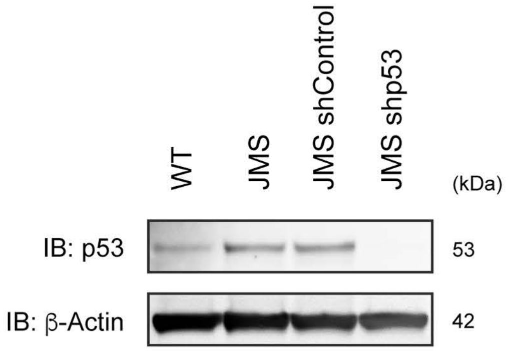


**Figure S5**
